# Cultural use of tropane alkaloids of *Brugmansia* species and cultivars in Colombia depends on local plant growth environment rather than genetic diversity

**DOI:** 10.1101/2023.11.02.565279

**Authors:** Sofia Rojas, Santiago Madriñan, Mark Stahl, Klaus Harter

**Affiliations:** Laboratorio de Botánica y Sistemáatica, Universidad de Los Andes, Bogotá, Colombia; Center for Plant Molecular Biology (ZMBP), University of Tübingen, 72076 Tübingen, Germany

**Keywords:** *Brugmansia*, secondary metabolites, tropane alkaloids, Andes of southern Colombia, traditional medicine, LC-MS

## Abstract

Borracheros or Angel Trumpets (*Brugmansia* spp.) of the *Solanaceae* plant family are known for producing scopolamine and other tropane alkaloids and are widely used by indigenous communities in the Andes for medicinal and ritual purposes. Growing different varieties has increased the popularity of *Brugmansia* species in recent years, leading to the promotion of hybridization to get new morphotypes. However, so far no medical or cultural use of plant extracts could be linked to selected *Brugmansia* species or cultivars. Moreover, until now there is only limited knowledge about the effect of hybridization on tropane alkaloid contents and/or diversity of *Brugmansia* species or cultivars. Here, a liquid chromatography-mass spectrometry (LC-MS) approach was applied to identify and characterize secondary metabolite contents in various *Brugmansia* species and hybrids from different geographic origins in Colombia. The analysis of the metabolic data sets revealed that tropane alkaloids and other secondary metabolites accumulate in a leaf and flower specific manner. Although some minor differences were found between *Brugmansia* subgroups, the only significant difference exists between plants collected in the Sibundoy Valley, where they have been domesticated by indigenous communities since a long time and those, which so far have not been used by humans at all. The different genetic constitutions of *Brugmancia* species and cultivars are not the dominating reason for their differences in tropane alkaloid contents and medical and cultural usages, as there is no clear taxonomical signature based on the metabolite properties for species segregation. Instead, their use by indigenous communities during ceremonies and for medical treatment such as wound healing, infection control, rheumatism and arthritis is rather caused by the plasticity of the individual plants to react to the local abiotic and biotic environments by the variation of secondary metabolite composition and by the differences in the bioavailability of the active compounds as result of the extraction method.

## 1. Introduction

In addition to other functions, plant secondary metabolites are related to plant-herbivore and plant-pathogen interactions, where the different chemical compounds, such as alkaloids, usually work in defence mechanisms (McKey, 1973; Wink 2008). However, recent studies have shown that the functions of secondary metabolites in plant biotic interactions are broader than thought before. For instance, secondary metabolites in root exudates shape microbiota in the rhizosphere in order to obtain better growth and defence conditions for the plants (Hu et al. 2018). Secondary metabolites also act as allelopathic compounds in plant-plant interactions (Bais et al. 2014). The synthesis of secondary compounds is related to the genetic repertoire and evolutionary history of the plant species as well as to the growth conditions of the individual plant (Pichersky and Gang, 2000; Wink, 2008; Ullrich et al. 2007).

Humans have been using plant secondary metabolites throughout history; from spices and herbs, to medicine and entheogens, among others (Simpson and Ogorzaly, 2013). Currently, approximately 25% of all medicinal drugs worldwide come from plant origin (Rates, 2001). Most of the pharmaceutical active compounds were discovered thanks to their initial usage by traditional communities, as e.g. Ayurvedic medicine in India or indigenous communities in the Americas (Schultes & Raffauf, 1990; Rates, 2001; Rätsch, 2005). One example of how secondary metabolites are used in medicine is *Brugmansia* Pers., a *Solanaceae* genus. The species are perennial small trees distributed in subtropical regions of South America, especially throughout the Andes and the south-eastern Amazon in Brazil (Hay et al. 2012). The plants are characterized by their pendulous and trumpet-shaped flowers, with long and thin corollas of various colours, varying from white, pink and pale yellow to bright yellow and red (Lockwood, 1973). Several authors have proposed different species for the *Brugmansia* L. (*B*.) genus, but no final consensus was reached so far. For this study, the Dupin and Smith (2018) taxonomy is used. Despite this ongoing debate, a consensus has been reached on the existence of two subgroups inside the genus: The *Sphaerocarpium* group, that consists of the high land “species” *B. sanguinea* and *B. arboreaa* and the *Brugmansia* group, that comprises the lower land “species” (*B. aurea, B. suaveolens, B. versicolor)* (Table S1). The difference between these subgroups is that their members are able to hybridize among each other within the identical subgroup but not with members of the other subgroup (Hay et al. 2012).

Among several names in different indigenous languages, *Brugmansia* (*B*.) is commonly known as Angel’s Trumpet or, in Spanish, as Borrachero, because of the shape of its flowers and the psychoactive effects of extracts derived from its tissues and organs. A huge variety of secondary metabolite tropane alkaloids (TA) were reported for all *Datura* ssp. and *B. aurea* as well as for *B. suaveolens* and two hybrids, varying mainly in the five subgroups of 3-monosubstituted tropanes, 3,6-disubstituted tropanes, 3-substituted-6,7-epoxytropanes, 3,6,7-trisubstituted tropanes and miscellaneous other alkaloids (Doncheva et al. 2006). A significant chemotaxonomic trait explained the differences between the *Datura* species with the presence or absence of entire TA subgroups. TAs are successfully used as chemotaxonomic fingerprints within *Solanaceae* to differentiate between tribes and genera (Griffin & Lin, 2000). In the *Solanoideae* clade, comprising *Brugmansia* spp. L. and *Datura* spp. L., a single evolutionary origin of the metabolic pathways that produce TAs was found, in which very high substrate-specific enzymes are active for every metabolic step (Wink, 2003; Boswell, 1999). Atropine, scopolamine and nor-hyoscine are the most abundant TAs in methanol extractions of *B. arborea* (Capasso et al. 2008). However, not all *Brugmansia* species haven been characterized chemically yet and until today no information is available on how the chemical compositions differ within a genus or whether there is any inheritance pattern.

Due to the presence of these TAs, *Brugmansia* species have been widely used for ritual purposes by several indigenous communities throughout the Andes (Schultes et al. 1982; Schultes & Raffauf, 1990; Capasso et al. 2003). For example, a very detailed study by De Feo (2003), carried out in northern Peru, describes two main types of usage. On the one hand, boiling water brews or ethanol tinctures are applied to the skin for relieving muscular or joint aches, especially related to rheumatism and arthritis, healing swellings, infections, anthelmintic activity and as general local analgesic. On the other hand, traditional healers and shamans use plant extracts for therapeutic-divinatory purposes, where they claim to see the illness affecting a certain individual of their tribe, the true self of people, to induce vivid dreams and as a truth serum. Due to the high toxicity, only rarely these applications involve eating of plant parts or drinking of plant extracts (Taita Juan Mutumbajoy personal communication; Rätsch, 2005). Comparable studies in the Sibundoy Valley of Colombia, home of the Kamentsa people, and with the Kofan people of the Amazon revealed similar medicinal uses (Schultes and Raffauf, 1990; Rätsch, 2005). Furthermore, Schultes and Raffauf (1990) also reported that either smelling or drinking extracts of these plants work as stimulant when people are tired or as promotors of visions and hallucinations. According to these studies, some *Brugmansia* varieties have specific purposes and ways of preparation, primarily selected by the colour of the flower (De Feo, 2003; Rätsch, 2005). This might suggest, that different TA profiles of *Brugmansia* subgroups and their hybrids might be the background for their differential human use. In this study, we aim to understand this in more detail with a focus on how hybridization might be influencing the phytochemical profiles that potentially led to the different medicinal and therapeutic-divinatory applications in traditional communities in Southern Colombia. In addition we try to reveal to what extend the local environment of individual plants, rather than their genetic background, contribute to different alkaloid contents and how different extraction protocols of local communities influence secondary metabolite contents and thus their medical use.

## 2. Results

### 2.1. Collection of plant samples

Leaves and flowers of 76 individuals were collected at 7 geographical places of Colombia: Inzá and Totoró - Cauca, where rural and indigenous communities co-inhabit; Sibundoy Valley - Putumayo, where the Kamentsa and Inga communities live; Bitaco - Valle del Cauca, private collection of *Brugmansia* (*B*.) with varieties from all northern South American low lands species, Garzón-Huila and in La Calera and Bogotá-Cundinamarca (Figure 1). Several individuals of the 5 species and several hybrids from both subgroups were collected, except for *B. arborea* which counts with only one individual as it occurs mostly in Ecuador.

**Figure 1.**
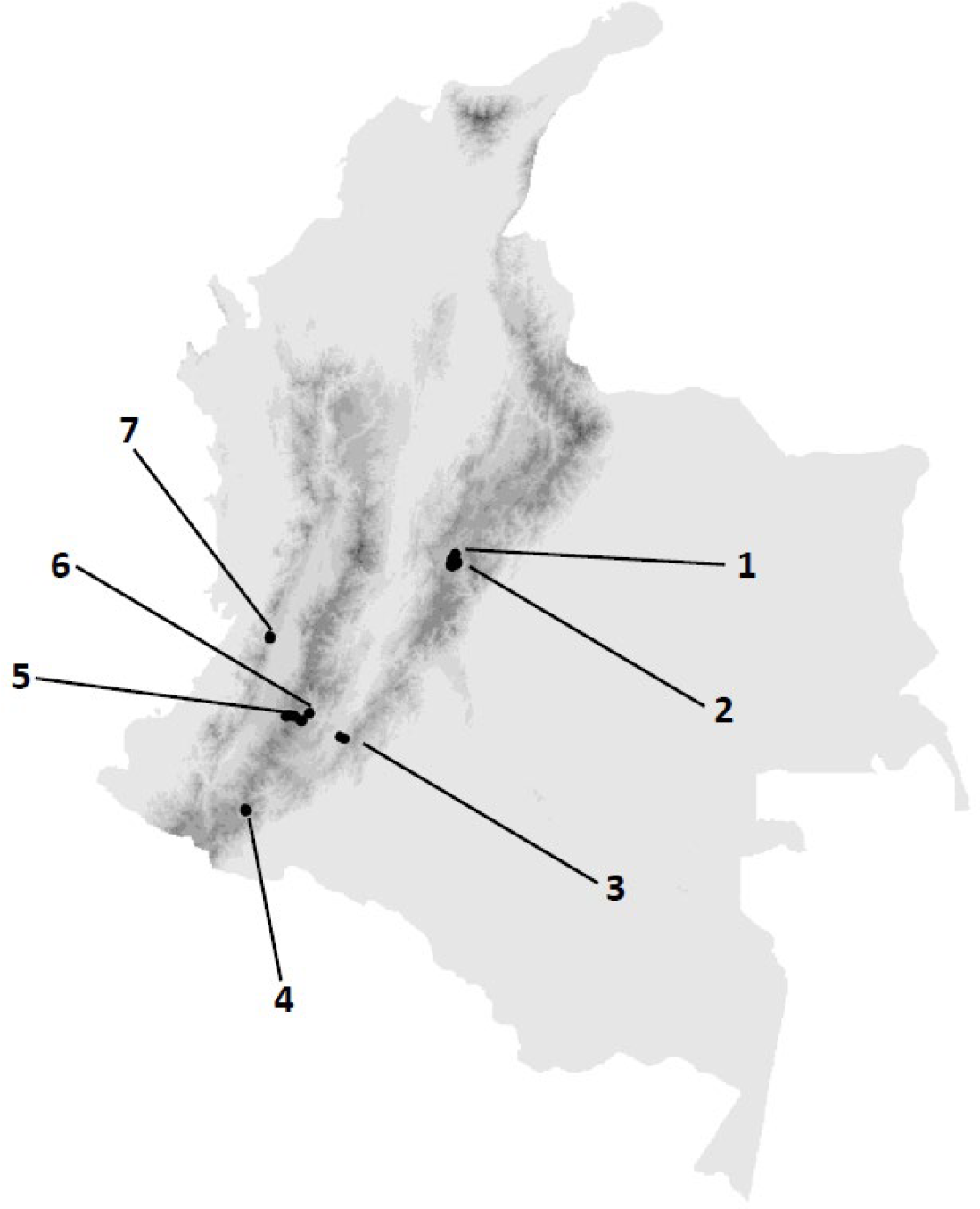
Map of the collection sites for the analysed *Brugmansia* samples on the Colombian territory. Each black dot is a GPS location of a specimen collected in the geopolitical locations of La Calera in Cundinamarca (1) and Bogotá (2), Garzón in Huila (3), Sibundoy in Putumayo (4), Inzá and Totoró in Cauca (5), (6) and Bitaco in Valle del Cauca (7).

### 2.2. Overall metabolic differences

After a methanol/water extraction of 76 leaf and flower samples we first applied a non-targeted LC-MS analysis in order to detect metabolic differences between all plant samples. We were able to detect a total of 27,836 m/z/ retention time pairs in positive and negative ESI modes. After an elaborate statistical analysis (see Material and Methods for details) 76 m/z/ retention time pairs were identified as potential differences of interest. Structural assignments resulted in nine putatively assigned compounds - four in positive and five in negative ESI mode (Table 1).

**Table 1.**
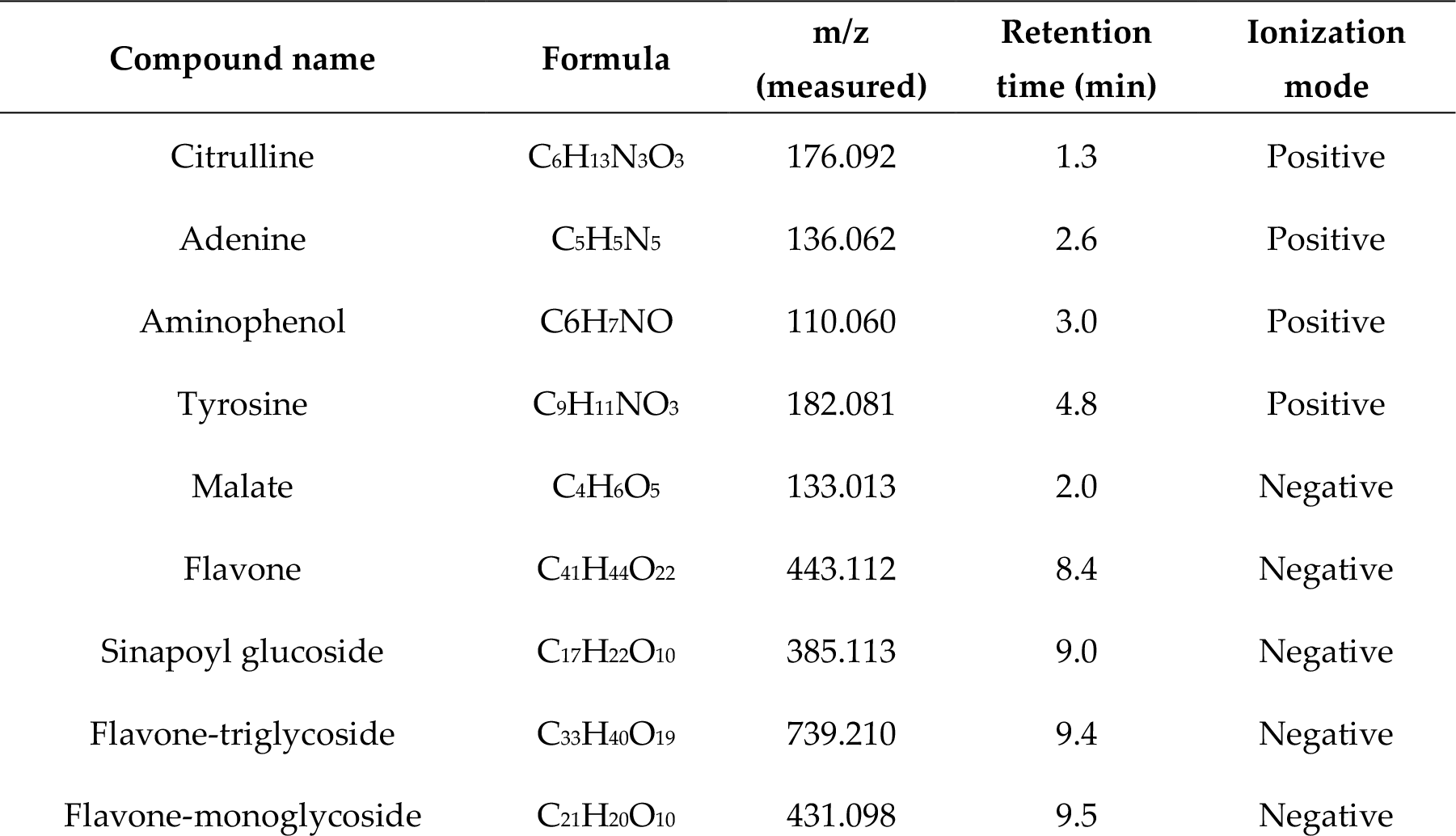
Metabolites from leaf and flower samples of *Brugmansia* species and hybrids after untargeted LC-MS analysis and non-target filtering processes.

The results of this approach revealed no clear differences in secondary metabolite composition between the *Brugmansia* species analysed, suggesting no or only little genetic differences in this respect.

### 2.2. Determination of tropane alkaloid (TA) contents with respect to plant organs, taxonomic classification and geographic origins

In order to search for TAs reported for the sister genus *Datura* by Doncheva et al. (2006), a targeted LC-MS approach was performed using the collected leaf and flower samples. On the basis of available reference standards and literature (Doncheva et al. (2006)) we were able to putatively identify 64 secondary metabolites with potential pharmaceutical effects (Table S2).

#### a) Distribution by plant organ

In the targeted approach, we first looked for metabolite distributions in leaf and flower samples. As shown in Figure 2, the leaves and flowers showed a taxonomic subgroup-independent, highly distinct profile (*p*-value < 0.001) demonstrating an organ-specific metabolite accumulation.

**Figure 2.**
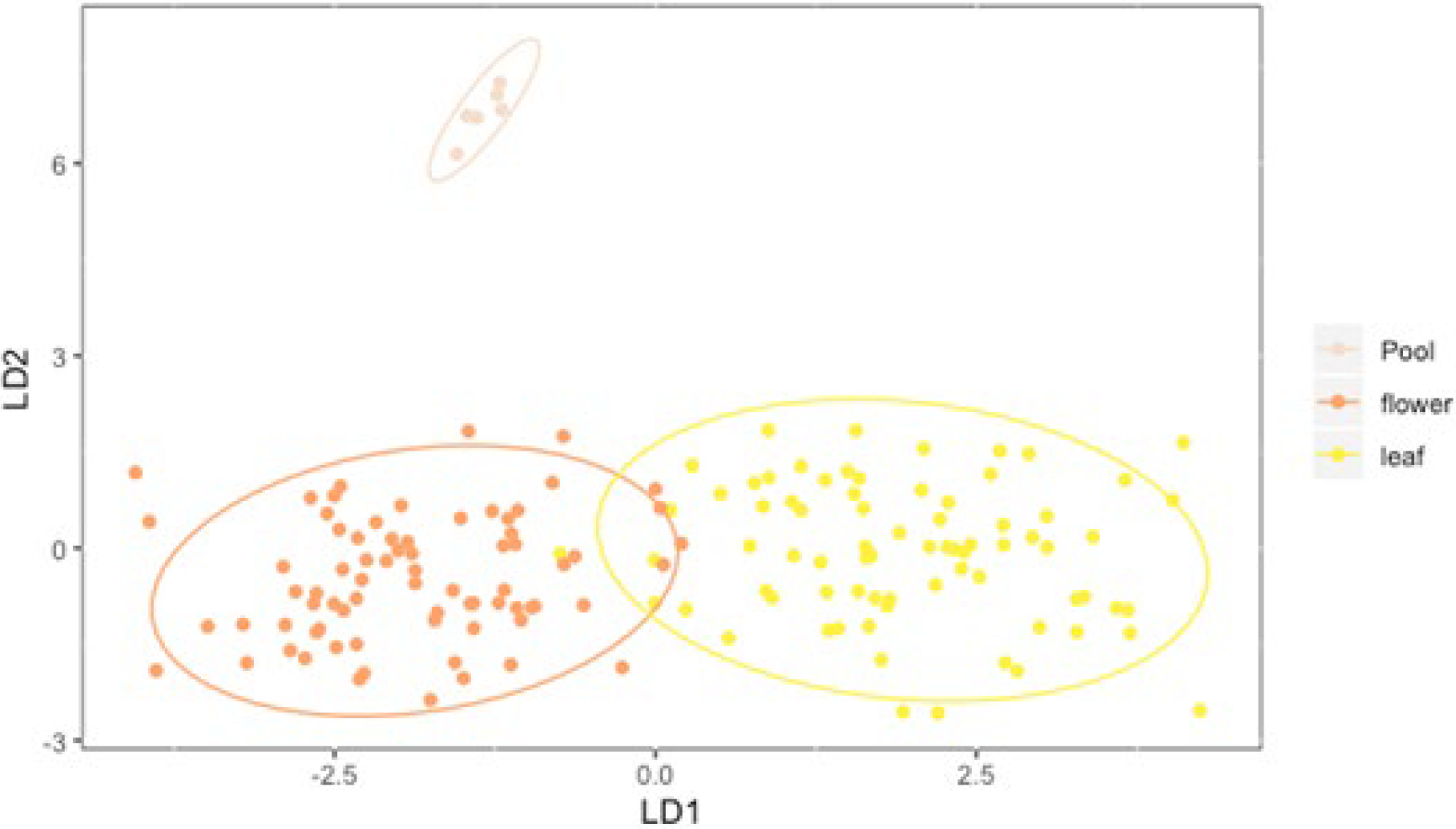
Metabolic compositions differ between leaves and flowers of the tested plant species and hybrids. A discriminating factor analysis (DFA) was performed with the obtained principle component analyses (PCAs) with all 64 metabolic compounds grouped by organs. X and Y axes are Linear Discriminants (LD) composed by the 64 metabolic peaks as variables. The ellipses represent 95% confidence intervals. A pool sample was created with 5 μl extract of every sample in order to assess the quality of the analysis.

#### b) Distribution by species / genetic background

When the occurrence of the 64 metabolites were discriminated for the taxonomic subgroups, a highly significant difference in their profiles (p value < 0.001) was observed between the *Brugmansia* and *Sphaerocarpium* subgroups of the genus, indicating that they are metabolically distinguishable (Figure 3).

**Figure 3.**
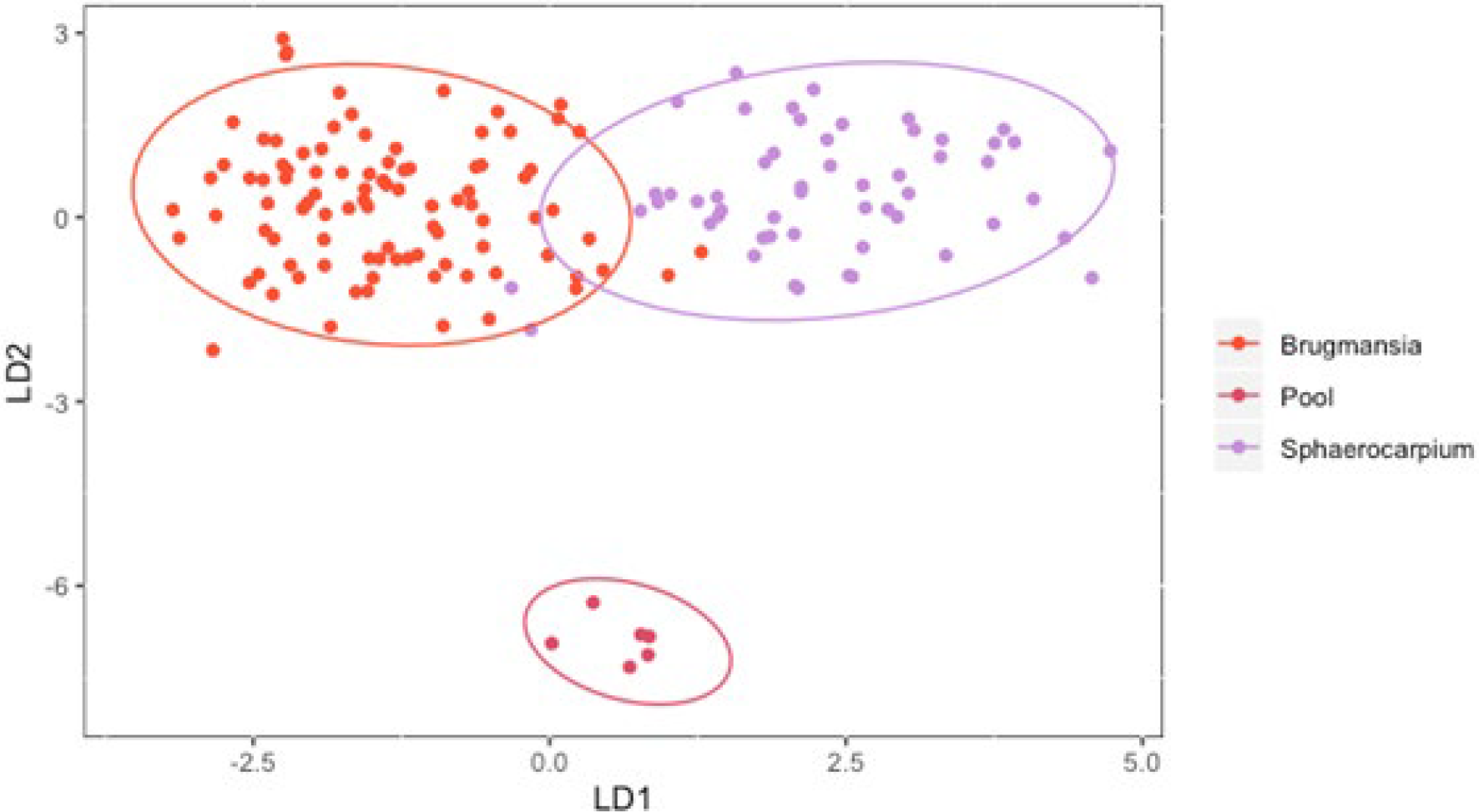
*Brugmansia* (Brugmansia) and *Sphaerocarpium* (Sphaerocarpium) subgroups are metabolically distinguishable. Discriminating factor analysis (DFA) performed with the obtained principle component analyses (PCAs) with all the 64 compounds by subgroups. X and Y axes are Linear Discriminants (LD) composed by the 64 metabolic peaks as variables. The ellipses represent 95% confidence intervals. The meaning of the “pool” is as in figure 2.

When the occurrence of the 64 metabolic peaks were discriminated by species, the hybrids of the *Brugmansia* subgroup were indistinguishable from the *B. versicolor* and *B. aurea* species, so were the hybrids of the *Sphaerocarpium* subgroup from *B. sanguinea* (*p*-value of Tukey test > 0.05) (Figure 4). However, there was a highly confident difference in the metabolite composition between the other species and the hybrid clusters (p-value < 0.006).

**Figure 4.**
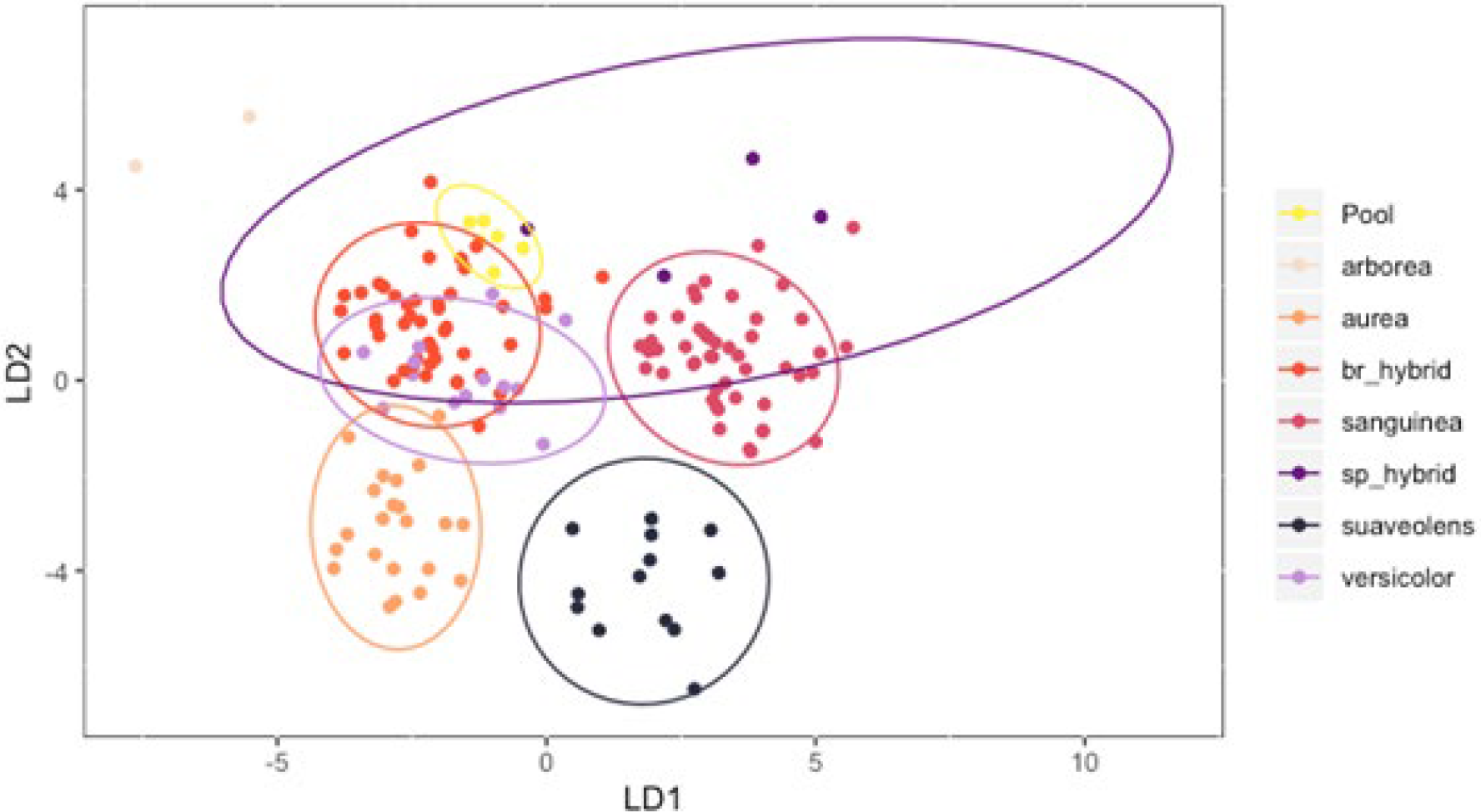
Differences but also similarities in the metabolite composition between the *Brugmansia* and *Sphaerocarpium* hybrids and species. Discriminating factor analysis (DFA) performed with the obtained principle component analyses (PCAs) with all the 64 metabolic peaks grouped by species and hybrids. The ellipses represent 95% confidence intervals. br_hybrid: all *Brugmansia* subgroup hybrids; sp_hybrid: all *Sphaerocarpium* subgroup hybrids. The meaning of the “pool” is as in figure 2.

#### c) Distribution by location

Next we were interested in the discrimination of metabolite occurrence in the plants by geographic location. As shown in Figure 5, the *Brugmansia* plants coming from the Sibundoy Valley were significantly distinct in their metabolic profile compared to those of the other regions (p value < 0.001). Moreover, two distinct subclusters were found in LD2, one representing the plants from Totoró and Inzá, both in the Cauca area, and the other representing the plants from the Bogotá, La Calera and Bitaco areas (Figure 5). The metabolic profiles of the plants from Garzón in the Huila region were somehow intermediate between the other two subclusters (Figure 5). In summary, there is a certain degree of differences in the metabolic composition in plants of different geographic origin.

**Figure 5.**
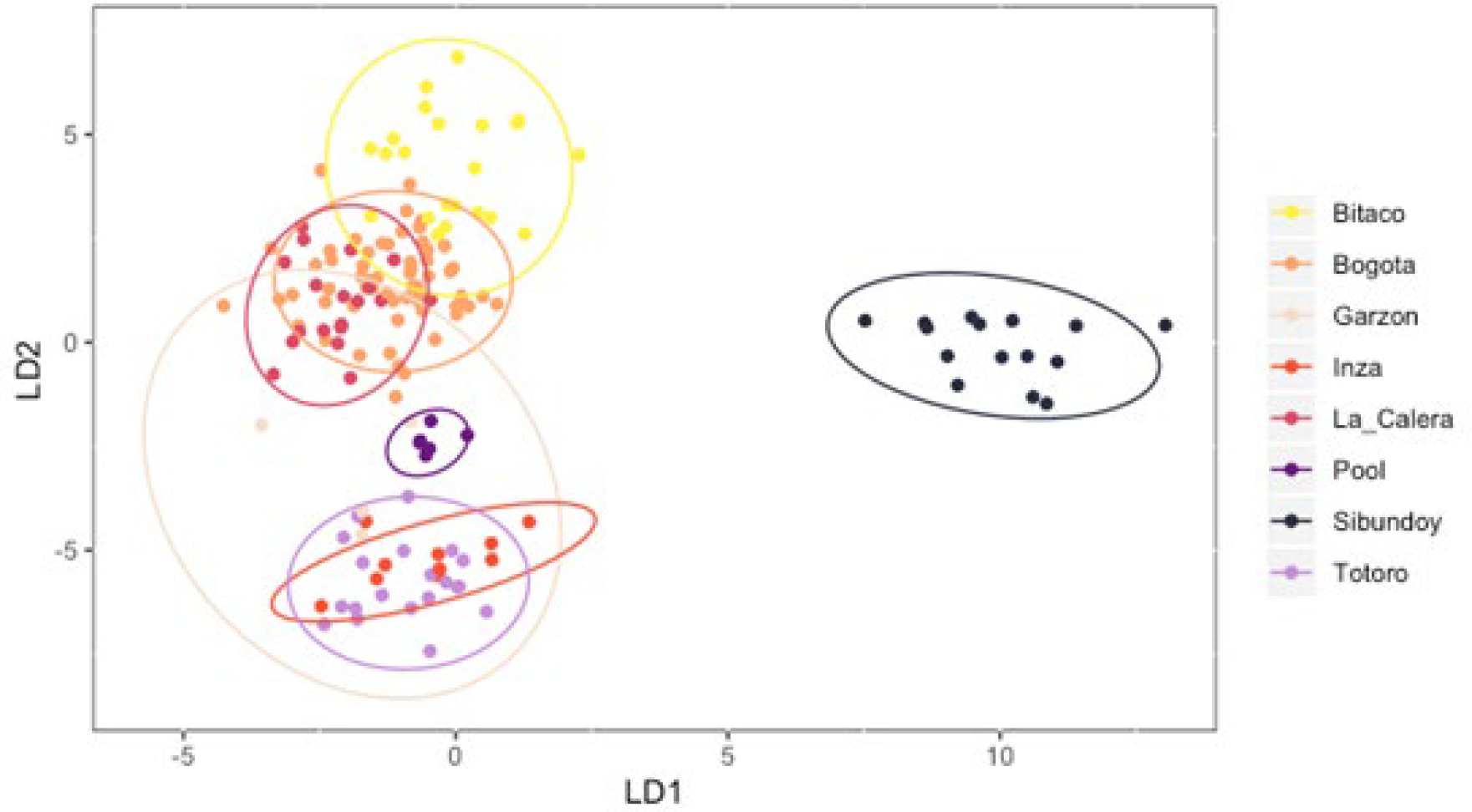
Metabolic compositions of the plants partially depend on their geographic origin. Discriminating factor analysis (DFA) performed with the obtained principle component analyses (PCAs) with all the 64 metabolic peaks grouped by geographical places. The ellipses represent 95% confidence intervals. The meaning of the “pool” is as in figure 2.

### 2.3. Diversity of minor TA in different Brugmansia species and hybrids

We identified scopolamine being the major TA in our samples. In terms of minor TAs, 4 different subclasses were found in the dataset: (i) TAs mono-substituted at positions 3 or 6, (ii) 3,6-disubstituted TAs, (iii) 3,6,7-trisubstituted TAs and (iv) 3-substituted-6,7 epoxy TAs (Table S2). As shown in figure 6, mono-substituted tropanes and 3,6,7-trisubstituted tropanes were the most abundant minor TAs in all species and hybrids. In addition, each *Brugmansia* species had its unique composition of minor TAs (Figure 6, 7).

**Figure 6.**
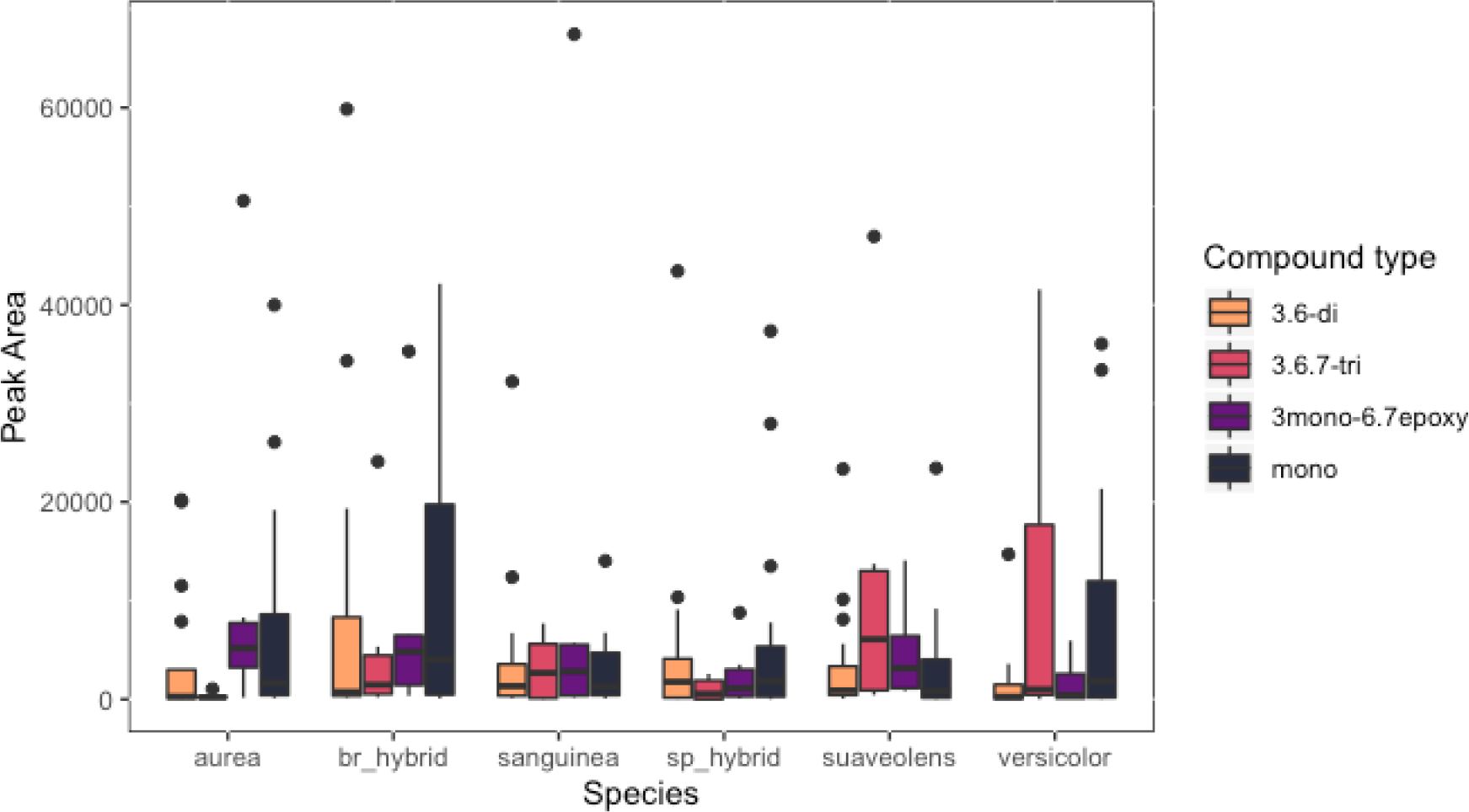
*Brugmansia* species and hybrids of different geographic locations are distinct in their general content of minor TAs. The average selected ion monitoring (SIM) peak area for each of the indicated substituted minor TA type is shown. The black bars represent the 95% confidence intervals and the black dots represent individuals measurements with TA-concentrations larger than the confidence interval. *For B. aurea* 11, *Brugmansia* hybrids 23, *B. sanguinea* 24, *Shpaerocarpium* hybrids 4, *B. suaveolens* 7 and *B. versicolor* 7 individuals were analysed.

**Figure 7.**
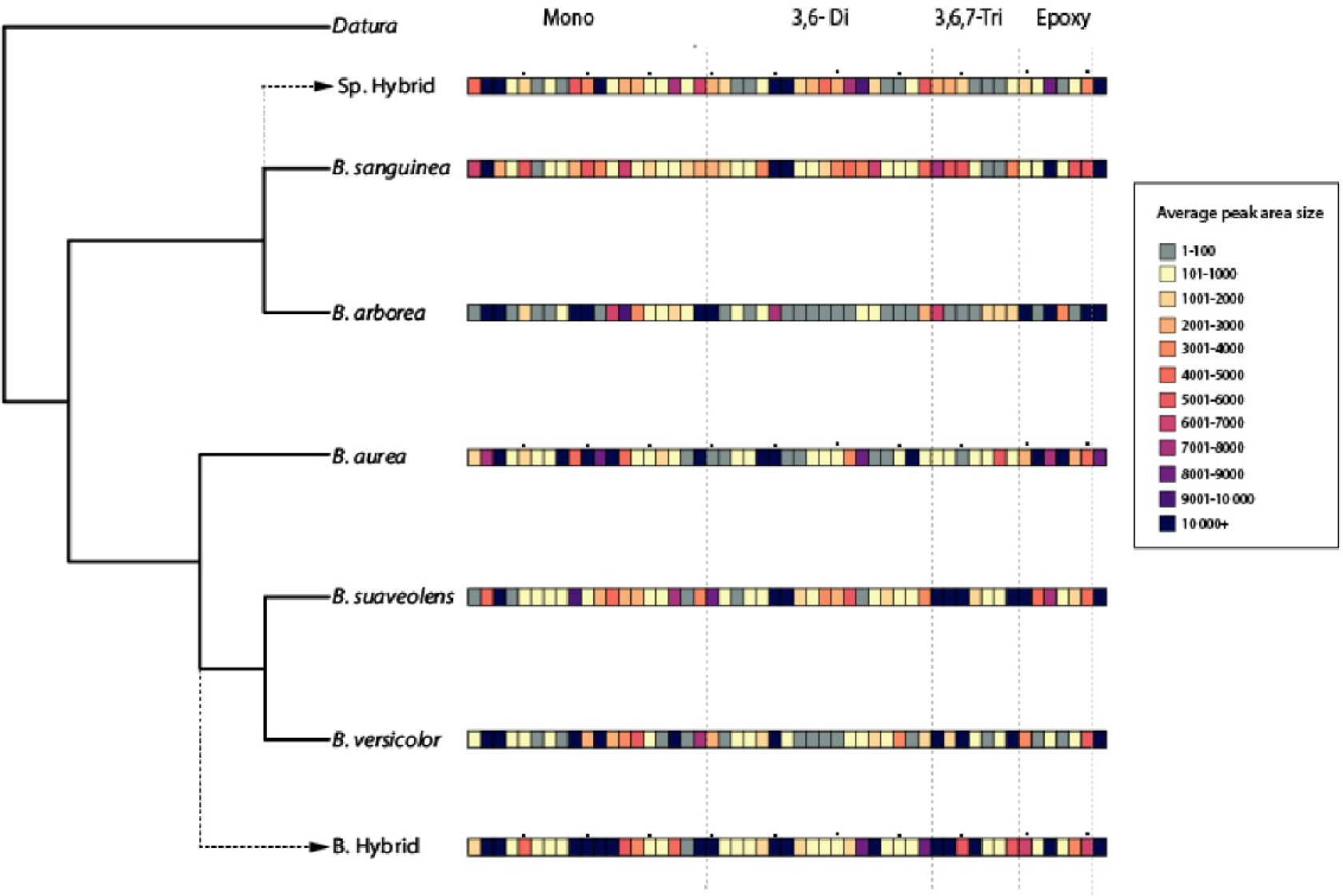
The *Brugmansia* species and hybrids show differences in the amount and composition of minor TAs. The average peak areas of all minor TAs per species or hybrids are shown colour-coded. The phylogeny was modified from Dupin and Smith (2018), adding the two sets of hybrids near the subgroup to which they belong. Each square represents a minor TA. The types of minor TAs are separated by the black dotted lines (mono: mono-substituted, 3,6-Di: 3, 6-di-substituted, 3,6,7-Tri: 3,6,7 –tri-substituted, Epoxy: 3-substituted-6,7-epoxy). The last square left of the dotted lines represent the average peak area for each type of minor TAs. For easier perception, the small dots above the squares are located every fifth square. The grey squares represent peak areas smaller than 100, which usually is background noise.

However, the TA variation within the samples of a given species from different geographic locations was often higher than the TA variation between the different species (Figure 6). This was particularly noticeable for the individuals of the *Brugmansia* hybrids and *B. versicolor*, where minor TAs varied more within the individuals of the hybrid/species from different regions than compared to the other species. Moreover, the *Brugmansia* hybrids had the highest content of minor TAs, whereas the *Sphaerocarpium* hybrids had the lowest one compared to the other species (Figure 6). In many cases, the samples of the hybrids did lack minor TAs that at least one of the parental species lacked as well (Figure 7).

### 2.4. TA diversity in Brugmansia samples obtained by different extraction methods

The indigenous communities use different methods to extract TAs and other secondary metabolites from *Brugmansia* species. To simulate these proceedings, we extracted both, leaf and flower samples, either in ethanol or boiling water and analysed the concentrations of the two major TA compounds scopolamine and atropine. However, we observed no clear extraction method - dependent differences in the concentrations of these two major TAs (Figure 8a, b).

**Figure 8.**
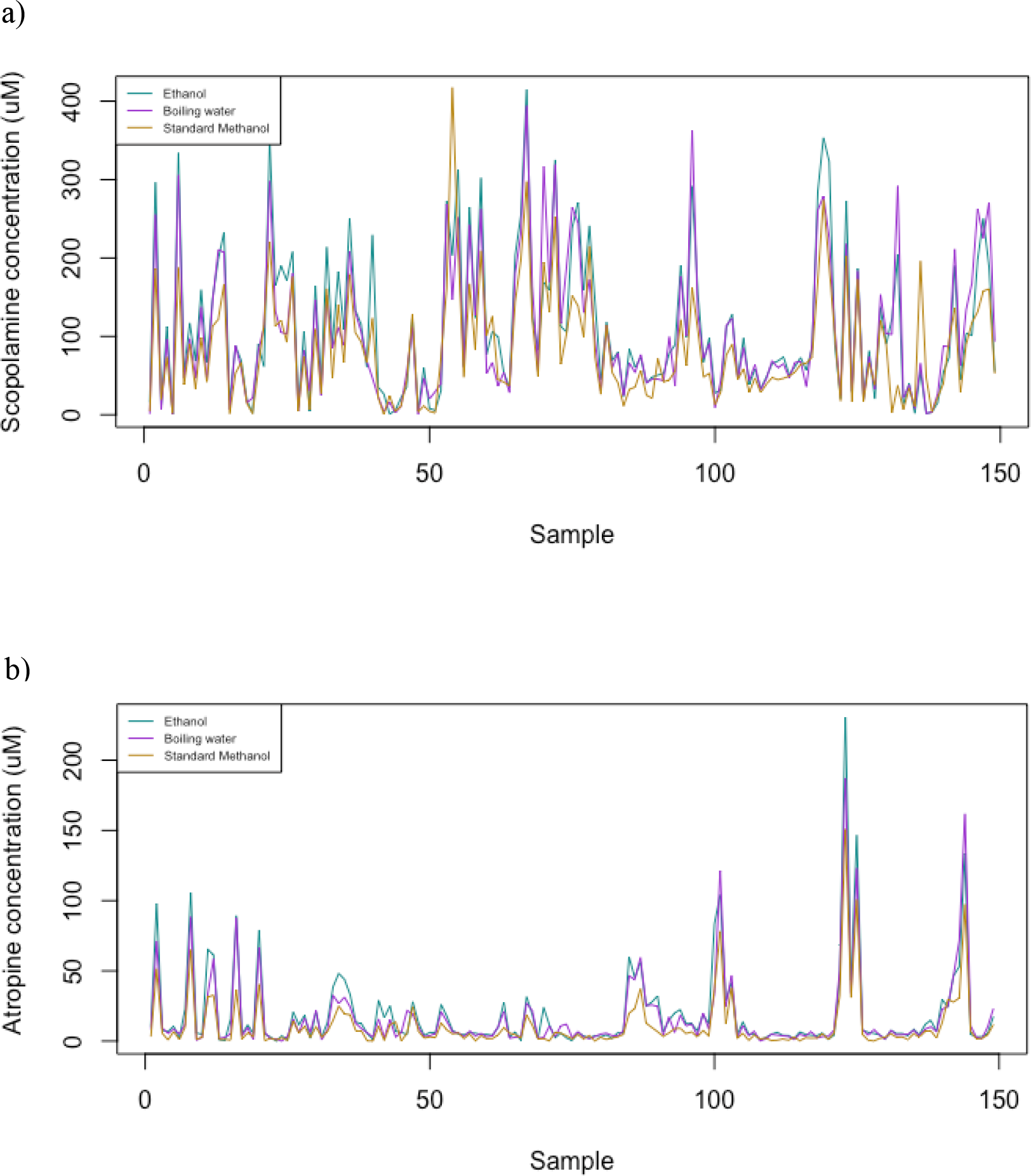
The concentrations of the two major *Brugmansia* TAs, scopolamine (a) and atropine (b), in the plant samples does not depend on the extraction method. The blue line represents the concentration of the TAs after ethanol extraction, the purple one after extraction in boiling water and the yellow one after the standard methanol/water extraction.

This suggests that the different medical applications are not based on different concentrations of the major TAs scopolamine and atropine while using different plant extraction methods.

To reveal the influence of the extraction methods on the composition of the minor TAs as well, the 64 metabolites addressed in this study were quantified in the samples in addition to scopolamine and atropine. When discriminating the results for the two extraction methods, *B. sanguinea* and the hybrids of the *Sphaerocarpium* subgroup were distinct from the other species and hybrids in LD1, whereas the species of the *Brugmansia* subgroup showed no significant differences (Figure 9a, b). When discriminating for the geographical origin of the plants, very similar metabolite profiles were observed after extraction of the samples with either ethanol or boiling water (Figure 9c, d). Once again, regardless of the species/hybrid, the differences in the plant samples from the Sibundoy valley compared to the other samples become obvious (Figure 9c, d).

**Figure 9.**
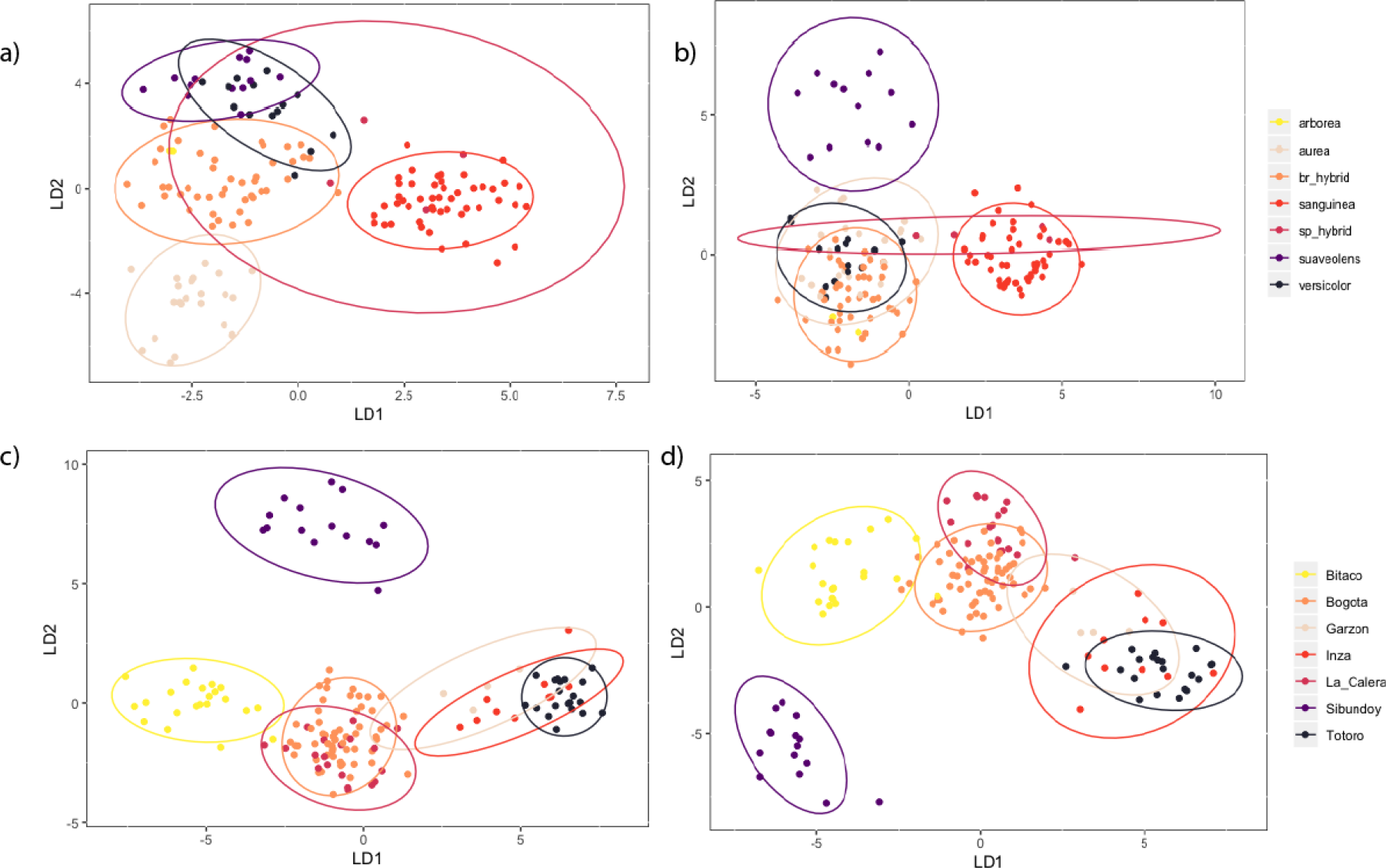
Ethanol and boiling water extractions show differences in how species were grouping with respect to species and geographic origin. Discriminating factor analysis (DFA) performed with the obtained principle component analyses (PCAs) with all the 64 metabolic peaks grouped by species after compound extraction in ethanol (a) and boiling water (b) extraction as well as grouped by geographic origin in ethanol (c) and boiling water (d). The ellipses are 95% confidence intervals.

Comparing figures 8 and 9, differences observed in the discriminating factor analysis (DFA) are rather caused by different concentrations of minor TAs, than those of scopolamine and atropine. In fact, each extraction method produced different contents for different minor TAs: For example, whilst 3,6-dihydroxytropane (m/z 158.12) was present in the boiling water extract, it was not present in the ethanol and standard methanol ones (Table S3). In addition, 19 compounds were found in the boiling-water and ethanol extracts only, but not in the standard methanol extracts, whereas 4 were only detected in the standard extracts (Table S3). This suggests that the sample extraction method does have an influence on the TA composition of the plant extracts, but it rather seems to fine tune TA compositions and thus the medical effect then being the major, discriminating parameter.

## 3. Discussion

In the present work, we analysed the compositions of TAs and further secondary metabolites in leaf and flower samples of the genus *Brugmansia* which were harvested at different geographical areas in Colombia. Generally, the main differences between sample groups were found in the compositions of the minor TAs rather than in the major TAs atropine and scopolamine. Different species have differential metabolite profiles, especially when comparing the members of the *Brugmansia* and *Sphaerocarpium* subgroups. As this corresponds with taxonomic grouping, it is most likely caused by slight genetic differences between the two species. The hybrids within the two subgroups are indistinguishable from their parental species with the exception of B. suaveolens in the *Brugmansia* subgroup, whose main morphological characteristics were not present in the hybrids. This latter observation may give interesting insights into the hybridization processes, where the two subgroups within the genus are starting to evolve not only morphological differences but also differences in their metabolome. This scenario is similar to the *Datura* genus, where the minor TAs have a better chemo-taxonomical potential for the taxonomy of the different species (Doncheva et al. 2006). As shown above, *Brugmansia* species have a high variety of mono-substituted tropanes, 3,6-disubstituted tropanes and 3-substituted-6,7-epoxytropanes. Although the species of the two different subgroups of the *Brugmansia* genus display general differences in the metabolic composition, the metabolite variation within the species is very high, especially with respect to the minor TAs. These differences might be, among others, due to the exposure of the individual plants to different climate and soil conditions, pathogene attacks, parasite and herbivore infestations, as it was described for several *Dubosia* species, another genus of the *Solanaceae* (Ullrich et al. 2007). This strongly suggests local environments as driving force for the metabolite patterns of the *Brugmansia* species.

Regarding the geographical origins, the *Brugmansia* samples from the Sibundoy Valley are strikingly different from those of the other regions, regardless of the applied extraction method. This is most likely because this *Brugmansia* population is under strong human selective pressure as the communities search for certain metabolic properties and promote varieties including hybrids that fulfil their expectations. This suggests a human made genetic shift in this case. The plants of the other regions are not subject to this high human selective pressure and, therefore, do not differentiate that strongly. This was also observed for other plant geni, subject to domestification in the Andean region, such as for lupine (*Lupinus mirabilis*; Atchinson et al. 2016).

In relation to leaf and flower samples, we observed a considerable difference in the metabolite profiles, regardless of the *Brugmansia* species. Our observation is in agreement with the results of a study by Alves et al. (2007) analysing individuals of *B. suaveolens* for their metabolite allocation and chemical dynamics throughout the plants life cycle (Alves et al. 2007). In this article Mickey (1974) and Rhoades (1979) are quoted for their theory of Optimal Defence, stating that the plant “defence should be preferentially allocated to the more valuable parts”. This theory is consistent with the chemical allocation of the main TA, scopolamine, during *B. suaveolens* growth and development. Initially, high amounts of scopolamine are found in the seed. With the progression of plant growth, scopolamine strongly accumulates in the meristem and young leaves. After the plant has reached the floral and fruit stages, the highest amount is detectable in these organs (Alves et al. 2007). This developmental and organ-specific accumulation of scopolamine is probably caused by the differential expression of the corresponding biosynthetic genes, as it was described for the TA biosynthetic pathway of *Hyoscyamus senecionis* (Dehghan et al. 2013). This developmental, organ-specific metabolite profile in leaf and flower tissues is most likely also one reason for the different usage and application of flower and leaf extracts by humans.

With respect to the extraction methods and solvents, there is some variation in the content of minor TAs. This might suggests that the different medical usages might be caused by differences in the minor TA contents. However, as the contents of the major TAs, scopolamine and atropine, show no significant differences, the minor TAs seem to rather modulate/fine tune the medical effects than being the major driving parameter. From a medical point of view, the route of application (oral or dermal) and the corresponding bio-availabilities and pharmacokinetic parameters of the pharmacologically active compounds, thus seem to have a much bigger influence on the medical effect than the detailed TA composition of the extracts. Several TAs found in *Brugmansia*, such as scopolamine, atropine and nor-hyoscine, were reported to act on various neuronal circuits, being able to block neuronal impulses. For instance, scopolamine is able to block the cholinergic system, involved in muscle contraction and withdrawal syndromes caused by discontinuation of opioid drugs and cocaine (Capasso & De Feo, 2003; Capasso et al. 2008; Bracci et al. 2013). Additionally, it was reported that extracts from *Brugmansia* species have an influence on the nociceptive response in rats, where the ability of detecting possible harm decreases (Muccillo-Baisch et al. 2010). Moreover, compounds of *Brugmansia* extracts are also able to interact with dopamine and serotonin receptors affecting human short term memory confirmation and the overall reward system in the brain, explaining their ritualistic use by indigenous communities (Nencini et al. 2006; Braccini et al. 2013). Anisodamine, another TA found in our *Brugmansia* extracts has the same effect on the central nerve system (CNS) than the main alkaloids mentioned above (Kohnen-Johannsen & Kayser, 2019). These impacts of TAs on the CNS do explain why extracts from *Brugmansia* species are used as pain killers and during ritual ceremonies. In combination with a high bio-availability and thus whole body, including CNS, exposure this suggests rather central nervous effects, whereas restricted, dermal applications with lower uptake rates and bio-availabilities will predominantly lead to topically limited, more local effects. This might very likely be the major reason for the very widespread and different medical applications of *Brugmansia* extracts in endogenous communities and should be investigated in future studies. In addition to the TA, we also identified phenolics, tannins, coumarins, flavones and many other secondary metabolites in our *Brugmansia* extracts (Table S2). These compounds are also associated with the treatment of rheumatic diseases and inflammations, but no compound group clearly correlated with the genetic background of the plants, its growth location, the extraction method or the medical use, suggesting that none of them is the major reason for a medical application.

In summary, the complex composition of TAs and other secondary metabolites in the extracts of *Brugmansia* plants and their profound impact on human health and wellbeing are the reason for their intense and continuous use in traditional medicine in the Americas. The range of medical applications is thereby extremely widespread, but correlates only very little with genetic background, taxonomy, geographic location, plant organ or extraction method. The TA and other secondary metabolite contents of a given plant individual seem to depend mainly on its individual local environment like e.g. light exposure, soil composition, pathogen and herbivore attacks on the one hand and its developmental stage. Moreover, for the differences in medical usage, the route of application and the different bio-availabilities (dermal versus oral, local versus whole body), seem to be much more important than the exact TA composition and/or content of the plant extracts.

## 4. Materials and Methods

### 4.1 Sample collection

Samples were collected in the departments of Putumayo, Cauca, Valle del Cauca, Huila and Cundinamarca, in Colombia from 07/2018 to 03/2019. Herbarium specimens were collected and stored at Herbario-ANDES and non-damaged leaves and flowers (when present) were collected in silica gel and afterwards lyophilized for 24 hours in order to preserve the samples until metabolomic assays could be carried out. All samples were taken in the mornings of the collection dates.

### 4.2 Standard metabolic extractions

10 mg of lyophilized samples of leaves and flowers were macerated independently with Qiagen® 5 mm stainless steel beads in a Retsch® Mill MM400 at a frequency of 30 collisions per second for one minute. 200 μl of an 80 % Methanol + 0.1 % formic acid + 9 μM Leu-Enkephalin solution (internal standard) were added to the powdered samples. The mixture was sonicated for 5 minutes followed by a 15 minute rest in ice. After a 10 minute centrifuge at 14000 rpm, the supernatant was transferred to a new tube and the same process was repeated with a 20 % Methanol + 0.1 % formic acid + 9 μM Leu-Enkephalin solution. Both supernatants were combined and centrifuged again. 5 μl of each sample was analysed by LC/MS. For absolute quantification external calibration of atropine and scopolamine was conducted from 500 nM to 100 μM each. In addition, a pool sample containing 10uL of each sample was prepared for the non-targeted analysis and served as a quality control during PCA analyses.

### 4.3 LC-MS

The alkaloid analyses were done with a Waters Acquity UPLC-SynaptG2 LC/MS system at the Center for Plant Molecular Biology (ZMBP) of the Universität Tübingen, Tübingen, Germany. The samples were ionized by positive and negative electrospray ionization (ESI) and the Q-TOF analyser was scanned from 50 to 2000 m/z at a scan rate of 0.5 s in MS and MSE mode in parallel. Chromatography was done at 30 °C with a 15 min gradient from 98 % water with 0.1 % formic acid to 99 % methanol with 0.1 % formic acid on a Waters Acquity C_18_ HSS T3 2.1 x 100 mm, 1.8 μm column.

### 4.4 Non-targeted analysis of compounds

Non targeted data analysis was done with ProgenesisQI v2.4 by Nonlinear Dynamics®. A pareto-scaled PCA was performed with all compounds detected and pairwise comparison between groups was done via OPLS-DA analyses. Highly relevant compounds evidenced in S-plots after the pairwise comparisons were tagged for further posterior evaluation and characterization. The resulting m/z retention time pairs and corresponding fragmentation spectra were searched in different databases (e.g. NIST, KEGG) for structural assignments.

### 4.5 Targeted analysis of secondary metabolites

The chromatograms were analyzed using Waters MassLynx software. All the m/z from the TA alkaloids reported in Doncheva et al. (2006) and some compounds found during the untargeted analysis were extracted from the TIC profiles using a ±0.01 Da chromatogram mass window and a ±0.3 min retention time window. The corresponding chromatographic SIM peak of each compound was afterwards integrated and in case of atropine and scopolamine calculated into absolute amounts by external calibration.

For further evaluation, in order to deal with missing data, a Random Forest imputation was carried out in MetImp 1.2 to the entire data set (Wei et al. 2018; Di Guida et al. 2016), followed by a normalization with a log function. A PCA was carried out in order to reduce redundancy and covariation between variables. The resulting PCs were then imputed into a discriminant factor analysis (DFA) in R in order to compare between subgroups within the genus, species, tissue type and locations, using psych (Revelle, 2018) and MASS (Venables & Ripley, 2002) R packages. Thanks to a report by Rätsch (2005) where coumarins are said to be present in *Brugmansia* species, different coumarins and coumarin-derivatives were looked for using Smyth et al. (2011) m/z values. Total and average peak areas of each TA subgroup were also compared between species and mapped on a Dupin & Smith. (2018) modified phylogeny.

### 4.6 Simulation of extraction according to the traditional medicinal applications

Two different traditional medicinal applications were simulated. Two sets of 10 mg of all lyophilized samples were weighted and afterwards macerated as described for the standard extraction. To simulated an ethanol extraction, 200 μl of an 80% ethanol + 0.1% formic acid + 9 μM Leu-Enkephalin (internal standard) solution was added to the first sample set. The samples were sonicated, left on ice and centrifuged as stated above. The second set of samples was extracted with 200 μl of distilled water (also containing 9 μM Leu-Enkephalin). The samples were sonicated, boiled for 20 minutes in a water bath, then left in ice and centrifuged as stated above. For external calibration of scopolamine, atropine, coumarin, p-coumaric acid and scopoletin standard material from 500 nM to 100 μM each were used. The analysis was performed as stated above. Absolute concentrations of scopolamine and atropine were calculated for each sample of the three extraction methods and compared between the different groups. For the boiling water and ethanol extractions, the concentration of coumarins, p-coumaric acid and scopoletin was quantified and compared in addition.

## Supplementary Materials

The following are available online at

## Author Contributions

Conceptualization, S.R. and S.M.; methodology, S.R., S.M. and M.S.; software, S.R. and M.S.; validation, M.S.; formal analysis, S.R. and M.S.; investigation, S.R., M.S. and K.H.; resources, K.H. and S.M.; data curation, S.R; writing—original draft preparation, S.R.; writing—review and editing, S.R., K.H. and M.S.; supervision, M.S.; project administration, K.H.; funding acquisition, K.H. and S.R.

## Acknowledgments

The authors would like to thank Analytics Facility of the ZMBP (University of Tübingen, Germany), especially Dr. Joachim Killian, Bettina Stadelhofer and Claudia Henner for their support in LC-MS and the *Botánica y Sistemática* and *Fisiología Vegetal* research groups at the Universidad de los Andes, Bogota, Colombia. We are also very grateful to the field assistants for their support: Juan Diego Castillo, Paula Velazquez, Sofía Medellín, Paula Cardozo, Pedro Luis Falla, Sandra Martinez and Juan Camilo Quiñones. In addition, the sample collection would have not been the same without Emilio Constantino’s and Alistair Hay’s private collections in Bitaco, Valle del Cauca, and without the Narvaez Muchavisoy and Mutumbajoy families in the Sibundoy Valley. Thanks to María Alejandra Meneses, Mateo Dávila and Lina Aragón for their comments on the text.

---

**Table S1.**
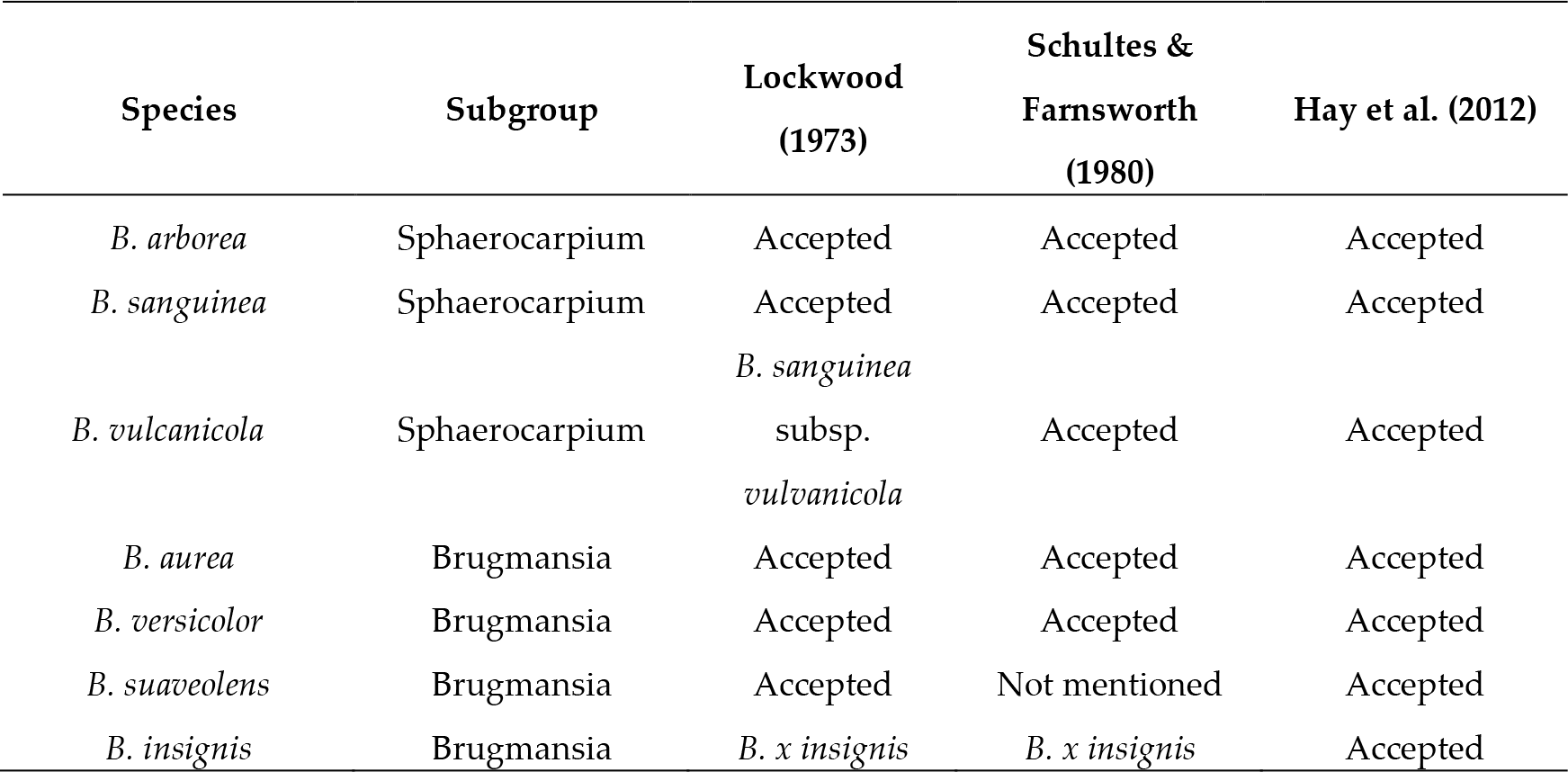
List and taxonomy of *Brugmansia* species used in this work according to different references

**Table S2.**
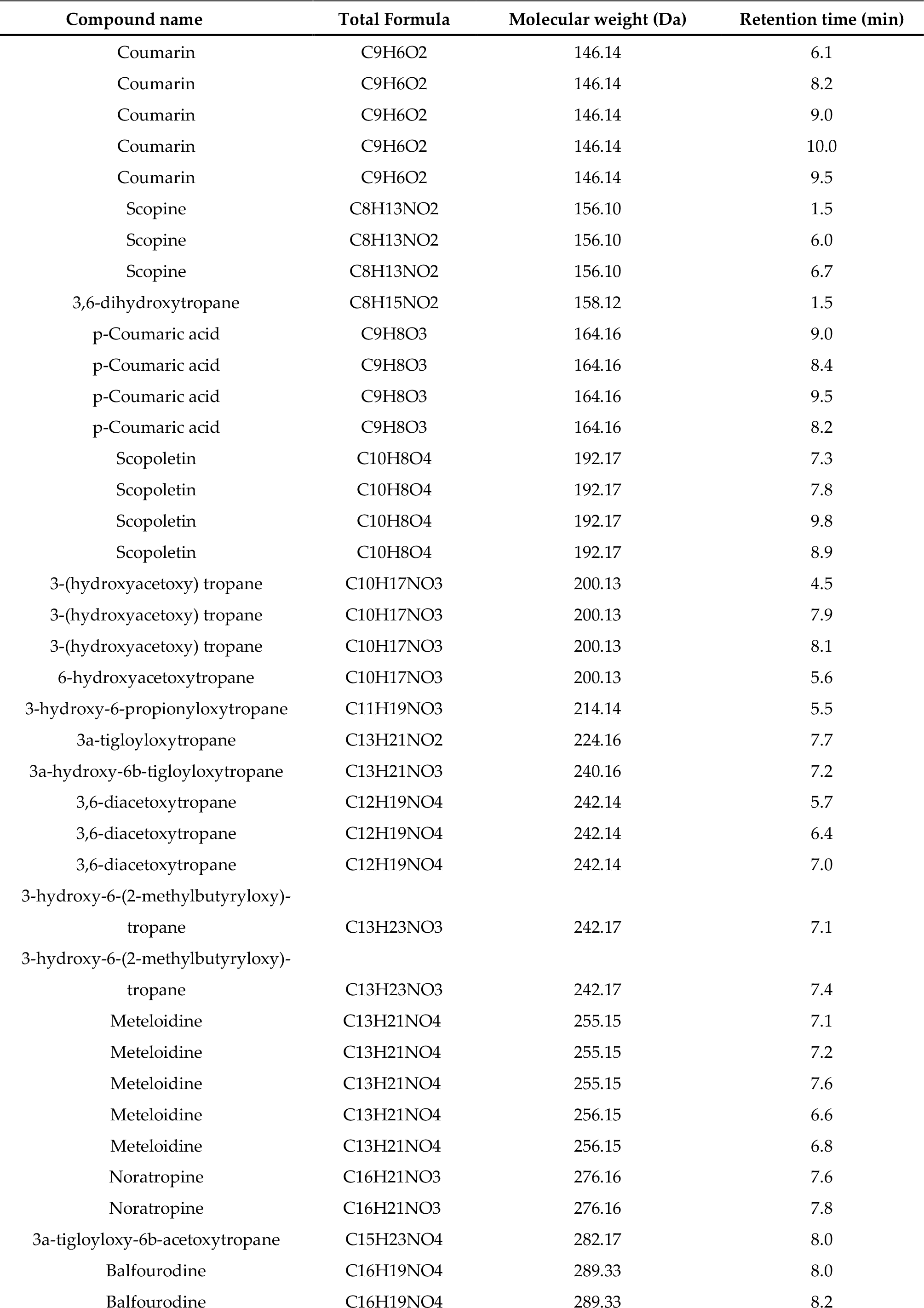
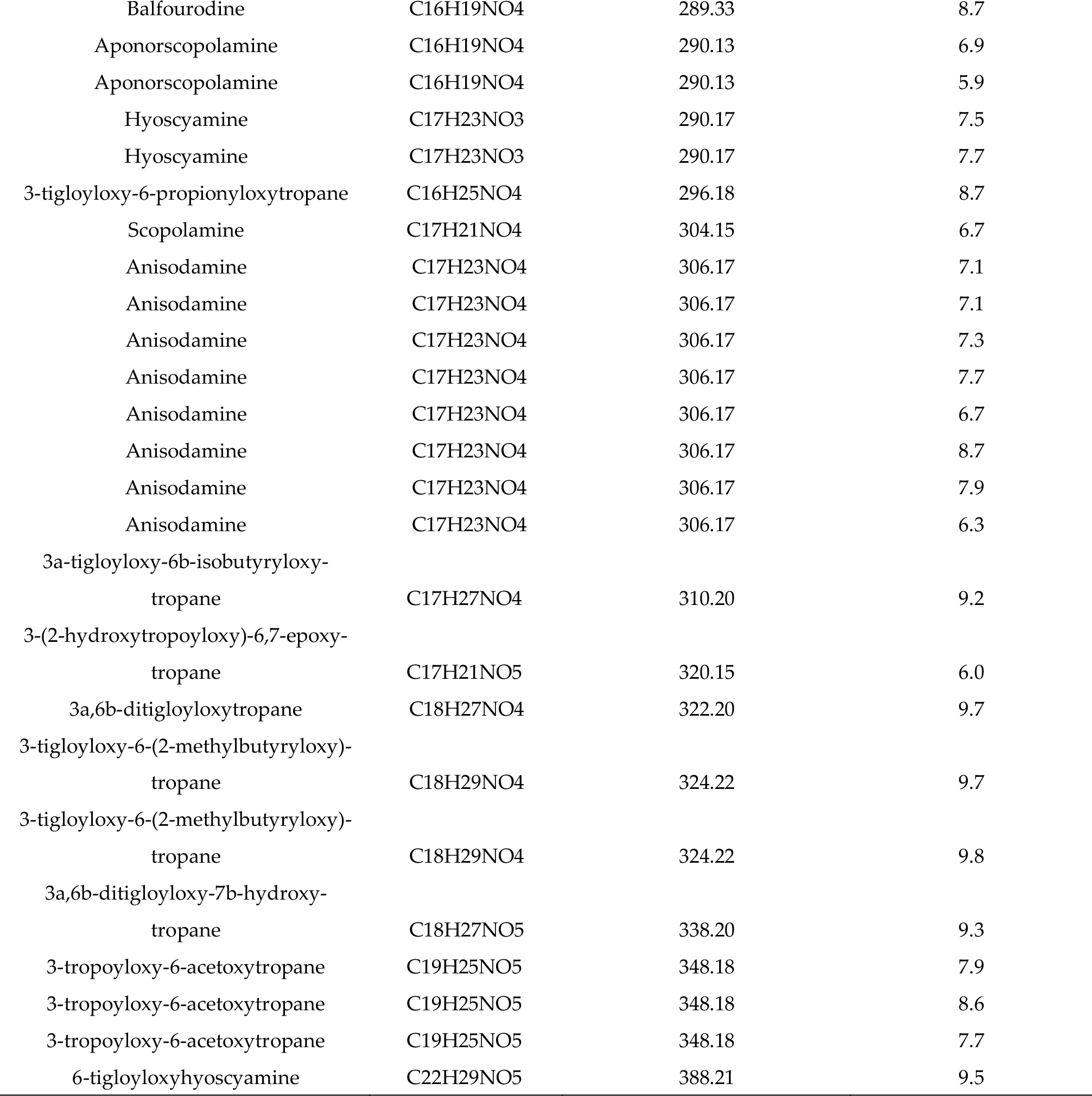
List of likely compounds identified in *Brugmansia* leaf and flower samples after standard extraction

**Table S3.** Differences in TA compositions (-absence or X presence) due to the differential extraction protocols. Presence of the compound in less than 10% of the samples were counted as absence. In type, Mono are monosubstituted tropanes; Di are 3,6-disubstituted tropanes; Tri are 3,6,7-trisubstituted tropanes and Epo are 3-monosubstituted -6,7 epoxy tropanes.

| Type | Compound | Retention time<br>(min) | Extraction Method |  |  |
| --- | --- | --- | --- | --- | --- |
|  |  |  | Standard | Ethanol | Boiling<br>water |
| Mono | 3-(hydroxyacetoxy) tropane | 2 | - | X | X |
| Mono | 3-(hydroxyacetoxy) tropane | 3.4 | - | X | X |
| Mono | Noratropine | 6.3 | - | X | X |
| Mono | Hyoscyamine | 7.5 | X | - | - |
| Mono | Apohyoscyamine | 8.06 | - | X | X |
| Mono | Apohyoscyamine | 9.8 | - | X | X |
| Di | 3,6-dihydroxytropane | 2 | X | X | - |
| Di | 3,6-dihydroxytropane | 3 | - | - | X |
| Di | 3,6-dihydroxytropane | 4.45 | - | X | X |
| Di | 3,6-dihydroxytropane | 7.5 | - | X | X |
| Di | 3-hydroxy -6- (2-methyl butyryloxy) -<br>tropane | 7.1 | X | - | X |
| Di | 3- hydroxy -6-propionyloxy tropane | 8.7 | - | X | X |
| Di | 3a- tigloyloxy -6b- isobutyryloxy tropane | 8.94 | - | X | X |
| Di | 3- tigloyloxy -6- (2-methylbutyryloxy)<br>tropane | 9.7 | X | - | - |
| Di | 3- tigloyloxy -6- (2-methylbutyryloxy)<br>tropane | 9.8 | X | - | - |
| Tri | Meteloidine | 6.46 | - | X | X |
| Epo | 3- (2-hydroxy tropoyloxy) -6,7- epoxy<br>tropane | 2 | - | X | X |
| Epo | 3- (2-hydroxy tropoyloxy) -6,7- epoxy<br>tropane | 5.36 | - | - | X |
| Epo | 3- (2-hydroxy tropoyloxy)-6,7- epoxy<br>tropane | 6.24 | - | X | X |
| Epo | 3- (2-hydroxy tropoyloxy)-6,7- epoxy<br>tropane | 6.5 | - | X | X |
| Epo | 3- (2-hydroxy tropoyloxy)-6,7- epoxy<br>tropane | 6.9 | - | X | X |
| Epo | Scopine | 6.24 | - | X | X |
| Epo | Aponorscopolamine | 6.9 | - | X | X |
| Epo | Aponorscopolamine | 7.74 | - | X | X |
| Epo | Scopoletin | 7.27 | X | - | - |
| Epo | Scopoletin | 8.73 | - | X | X |
| Epo | Scopoletin | 9.9 | - | X | X |

**Figure S1.**
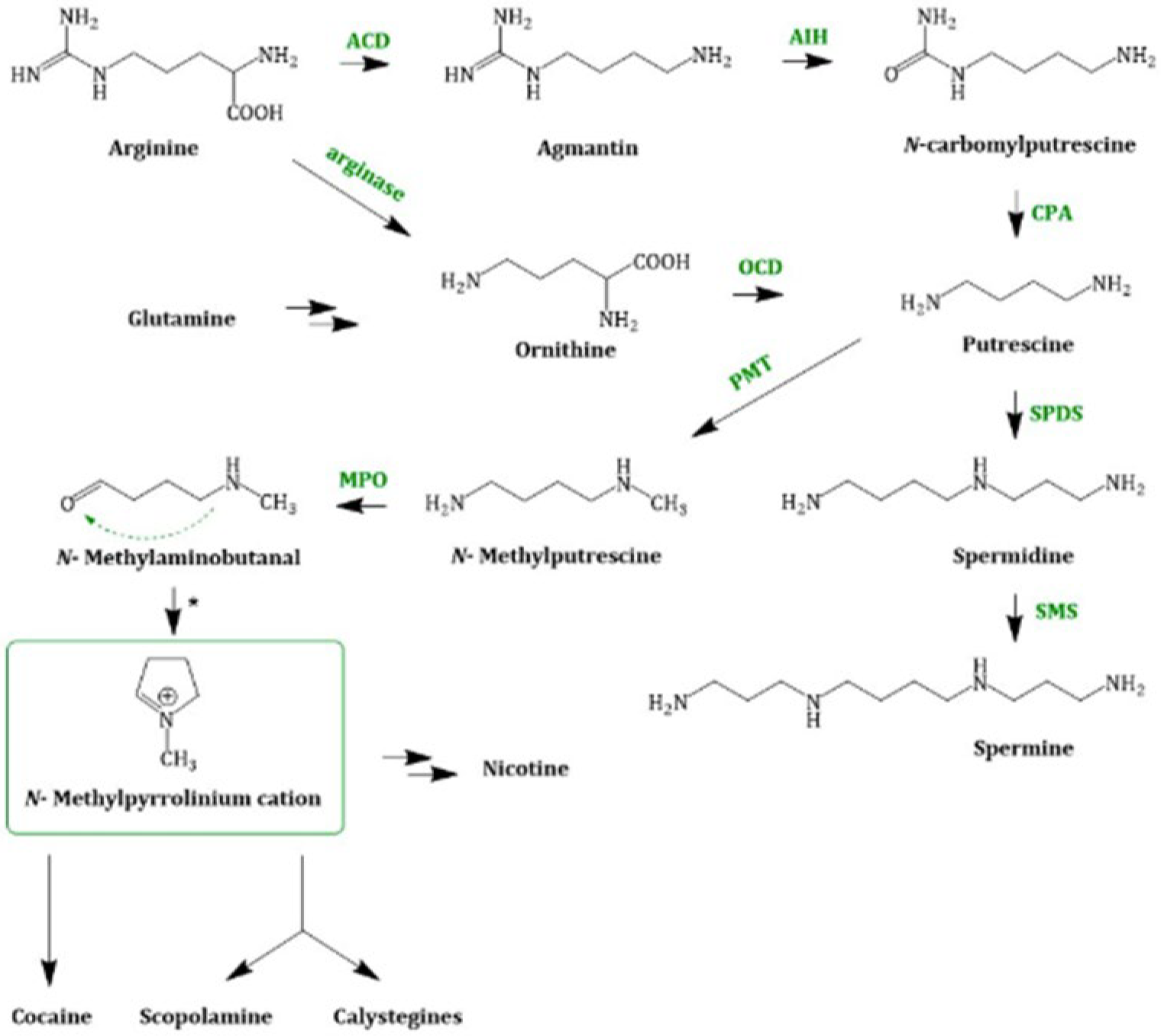
Schematic overview of the biosynthetic pathway of scopolamine according to Kohnen and Kayser (2019). Please note that arginine is also the precursor for other alkaloids such as cocaine and nicotine.

